# Primate Hippocampus Reveals Distinct Rules for Associative Synaptic Plasticity

**DOI:** 10.64898/2026.05.18.725835

**Authors:** Anoop Manakkadan, Krishna Kumar, Yee Song Chong, Wong Lik-Wei, Sheeja Navakkode, Yuk Peng Wong, Soong Tuck Wah, Camilo Libedinsky, Sreedharan Sajikumar

## Abstract

Long-term potentiation (LTP) is a key cellular mechanism underlying learning and memory, but its conservation across species remains unclear. Using nonhuman primates (NHPs), we examined hippocampal synaptic plasticity at Schaffer collateral– CA1 synapses. Theta-burst stimulation (TBS) reliably induced LTP in NHPs, comparable to rodents. However, unlike rodents, TBS in NHPs readily engaged synaptic tagging and capture (STC), indicating a lower threshold for associative plasticity. This was accompanied by increased expression of plasticity-related proteins, including PKMζ and BDNF, suggesting enhanced recruitment of protein synthesisdependent stabilization mechanisms. These findings reveal a species-specific divergence in the molecular regulation of persistent synaptic plasticity and identify an evolutionary specialization in mechanisms supporting associative memory. Together, our results highlight limitations of rodent models in fully capturing human-relevant memory processes and underscore the importance of primate systems for translational neuroscience.

## Introduction

Memory is widely regarded as an evolutionarily conserved cognitive function, although its complexity varies across species. At the cellular level, memory formation is thought to rely on activity-dependent changes in synaptic strength, commonly referred to as synaptic plasticity. However, the precise mechanisms by which neural circuits stabilize and adapt to these plastic changes to support complex learning remain poorly understood. Recent comparative studies between rodents and non-human primates (NHPs) provide compelling evidence for an evolutionarily conserved memory coding scheme, suggesting that fundamental aspects of information storage may be preserved across species (*1*). However, differences have also been uncovered between rodents and primates. For example, while the rodent hippocampus contains place cells that encode the animal’s location in space, the primate hippocampus contains view cells that encode the animal’s gaze direction(*2*).

Activity-dependent changes in synaptic strength, such as long-term potentiation (LTP), are widely proposed to underlie memory storage in the brains of mammals, including humans (*3, 4*). Although most studies of synaptic plasticity have been conducted in rodents, evidence from primate brains suggests that the fundamental mechanisms of LTP are conserved across species. Consistent with this, LTP can be reliably induced at both mossy fiber-CA3 and associational-commissural synapses in hippocampal slices from cynomolgus monkeys, with induction properties closely resembling those observed in rodent hippocampus (*5*). Similarly, NMDAR-dependent LTP has been reported in human cortical tissue (*6*) and in the primate visual cortex, where it exhibits Hebbian and associative properties comparable to those described in rodents (*7*). Moreover, in vivo studies in awake monkeys have demonstrated long-lasting spike-timing-dependent plasticity in the sensorimotor cortex, further supporting the presence of conserved synaptic plasticity mechanisms in primate brains (*8*). However, despite these similarities, it remains unclear whether differences exist in the mechanisms governing associative synaptic plasticity across species, an important question that warrants further investigation.

One of the most compelling aspects of LTP lies in the molecular mechanisms that support its persistence, a feature considered fundamental to memory storage. Importantly, LTP is not a unitary phenomenon; its expression varies depending on the specific synapse and neural circuits involved. Distinct patterns of stimulation can give rise to mechanistically different forms of LTP (*9*). Among these, theta burst stimulation (TBS) and high-frequency stimulation (HFS) are two commonly employed protocols for inducing LTP (*10*). Although both paradigms engage overlapping molecular targets, TBS-induced LTP (TBS-LTP) is particularly sensitive to various experimental manipulations, including aging (*11*), stress (*12*), serotonin modulation (*13*), endocannabinoid signaling (*14*), and adenosine receptor activity (*15*). This heightened sensitivity makes TBS-LTP a potentially more informative model for examining and comparing LTP mechanisms across species. Accordingly, TBS-LTP offers a physiologically relevant and experimentally advantageous platform for comparative analyses of synaptic plasticity across species.

TBS-LTP is particularly well suited for translational comparisons, as it is largely dependent on local protein synthesis, thereby minimizing nuclear involvement and emphasizing synapse-autonomous mechanisms. Moreover, LTP induced by TBS fails to engage associative plasticity mechanisms such as synaptic tagging and capture (STC) in rats (*16, 17*). STC refers to a process in which weakly activated synapses set transient molecular “tags” that allow them to capture plasticity-related proteins (PRPs) synthesized in response to a strong, temporally proximal stimulus, thereby stabilizing long-term synaptic changes (*3, 18, 19*). The inability of TBS-LTP to recruit STC may reflect insufficient availability of key PRPs at tagged synapses, such as PKMζ or a failure of TBS to elicit cell-wide protein synthesis and distribution necessary for associative synaptic consolidation (*16*).

Metaplasticity, the plasticity of synaptic plasticity, has been shown to modulate the mechanisms underlying STC through the recruitment of distinct PRPs (*16, 20*). By dynamically adjusting the threshold for LTP, metaplasticity establishes an optimal synaptic state for associative learning. Importantly, the LTP modification threshold is likely to vary across species, reflecting differences in the intrinsic complexity of neural circuits and regulatory processes. In support of this notion, comparative synaptic proteomic analyses of the hippocampus have revealed pronounced differences in protein expression profiles between rodents and primates (*21*). These observations raise the possibility that species-specific expression of distinct synaptic proteins may underlie functional differences in cellular associative properties across mammals.

In this study, we show that TBS reliably induces robust and persistent LTP lasting up to 6 hours in hippocampal slices from both rodents and NHPs. Strikingly, whereas TBS-induced LTP in rats failed to engage associative plasticity, the same stimulation paradigm facilitated STC in primates. Consistent with this divergence, TBS elicited higher expression levels of *BDNF* and *CaMKIV* transcripts in primate hippocampus compared with rats. At the protein level, Western blot analyses revealed a pronounced upregulation of PKCζ and BDNF following TBS-LTP in primates, an effect that was absent in rodent hippocampal slices and control NHP tissue. Pharmacological studies further uncovered species-specific mechanisms of LTP maintenance. In rats, BDNF was required for the persistence of LTP but did not contribute to its associative expression. In contrast, in primates, BDNF and PKMζ exhibited functional redundancy, such that inhibition of either molecule alone failed to disrupt LTP maintenance. However, simultaneous inhibition of both BDNF signaling and PKMζ activity abolished the late phase of LTP, revealing an interdependent molecular framework that sustains synaptic potentiation in the primate hippocampus.

## Results

### TBS-Induced LTP Promotes Synaptic Tagging and Capture in NHP Hippocampal Slices

To examine species specificity in the induction of TBS-LTP, we used hippocampal slices from rats and monkeys. Using a two-input experimental model (Fig. 1A i-iii), we show that a 5-Hz theta-burst stimulation (TBS) protocol reliably induced late-phase LTP (L-LTP) in both rat and NHP hippocampal slices. In rodents, this form of LTP has been shown to depend on protein translation but not transcription (*17*).

**Figure 1.**
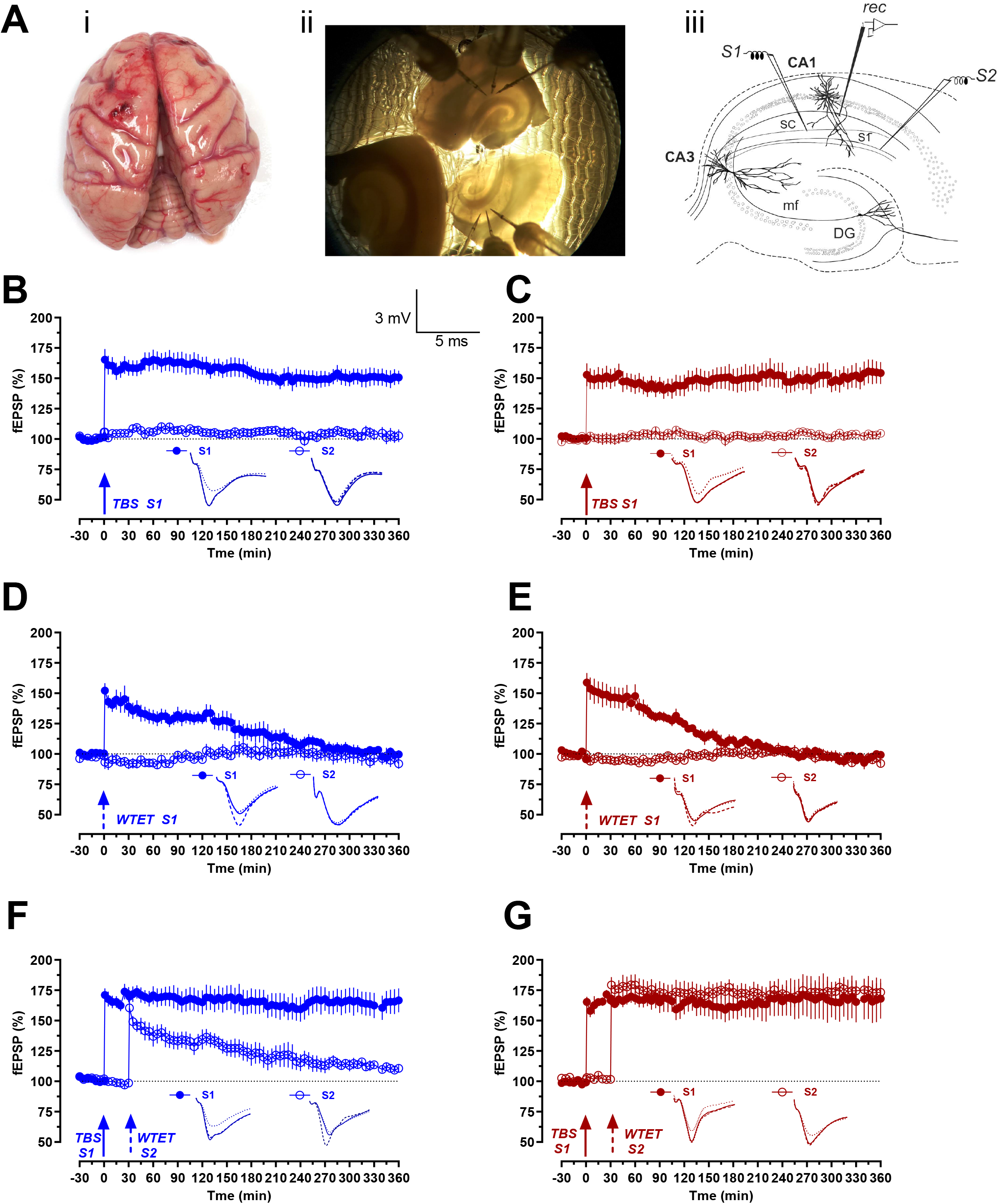
Species-specific differences in TBS-induced synaptic plasticity and synaptic tagging and capture (STC) in hippocampal CA1. **(A)** (i) Isolated NHP brain. (ii) Representative image of NHP hippocampal slice preparation with electrodes positioned in the CA1 region. (iii) Schematic of a hippocampal slice illustrating two independent synaptic inputs (S1 and S2) onto CA1 pyramidal neurons. Field excitatory postsynaptic potentials (fEPSPs) were recorded from apical dendrites in CA1 using a recording electrode (rec). (**B-C)** TBS-induced late-phase LTP (late-LTP). Following a stable 30 min baseline, theta-burst stimulation (TBS; 5 Hz) delivered to input S1 induced robust and persistent potentiation in both rat (B) and NHP (C) hippocampal slices. Potentiation was significant from 5 min post-TBS and persisted up to 6 h, whereas the control pathway S2 remained stable. **(D-E)** Early-LTP induced by weak tetanic stimulation (WTET). WTET applied to S1 induced transient potentiation in both rat (D) and NHP (E) hippocampal slices. Early-LTP decayed to baseline within ∼155-200 min, with no sustained potentiation observed in either species. **(F-G)** In rat hippocampal slices (F), TBS-induced late-LTP in S1 failed to support STC, as WTET-induced potentiation in S2 decayed to baseline, indicating lack of plasticity-related protein (PRP) capture. In contrast, in NHP hippocampal slices (G), WTET delivered to S2 30 min after TBS in S1 resulted in the conversion of early-LTP into persistent late-LTP, demonstrating robust STC expression. Both S1 and S2 pathways remained significantly potentiated throughout the 6 h recording period. In all panels, figures shown in blue represent experiments conducted in rats, whereas those in dark red represent experiments in nonhuman primates (NHPs). Solid and dotted arrows indicate the time points of application of TBS and WTET, respectively. Analog traces depict representative S1 and S2 fEPSPs: 30 min before any experimental manipulation (dotted line), 30 min (hatched line) and 360 min (solid line) after TBS or WTET. Vertical scale bar: vertical: 3 mV, horizontal scale bar: 5 ms.

As a control experiment, we first investigated TBS-LTP in rat hippocampal slices. After recording a stable baseline for 30 min, TBS-LTP was induced in synaptic input S1 (Fig. 1B, filled blue circles). Statistically significant potentiation was observed at Schaffer collateral-CA1 synapses in rats starting from the first recording after TBS, at 5 min (Wilcox test, *P* = 0.0313; U-test, *P* = 0.0022), and persisted until 6 h (Wilcox test, *P* = 0.0313; U-test, *P* = 0.0022), compared to its own baseline or the control input S2. Baseline responses recorded from synaptic input S2 remained stable (Fig. 1B, open blue circles).

Given the involvement of the primate hippocampus in higher-order cognition, we reasoned that neural activity patterns may differ between NHPs and rodents. Accordingly, we tested whether the NHP hippocampus exhibits similar thresholds for TBS-induced LTP. As in Fig. 1C, a stable baseline was recorded for 30 min in both S1 and S2 (filled and open brown circles), after which TBS was delivered to input S1. TBS induced a potentiation that was statistically significant from 5 min up to 6 h (Wilcox test, *P* = 0.0313; U-test, *P* = 0.0022). These results demonstrate that, similar to rats and mice, TBS can induce a persistent potentiation lasting up to 6 h in NHP hippocampal slices.

As activity-induced synaptic plasticity was comparable in rat and NHP hippocampal slices, we next asked whether short-term potentiation, such as early-LTP, is similarly preserved in NHPs. In rat hippocampal slices, early-LTP was induced using a WTET protocol (Fig. 1D). WTET elicited a transient potentiation that persisted only until 155 min relative to its own baseline (Wilcox test, *P* = 0.0313) and until 155 min when compared with the non-tetanized S2 pathway (U-test, *P* = 0.0095, Fig.1D, filled blue circles). To examine short-term synaptic plasticity in NHP hippocampal slices, WTET was next delivered to S1 (Fig. 1E). In this case, WTET induced an LTP that was significant only until 155 min relative to its own baseline (Wilcox test, *P* = 0.0313) and 155 min when compared with S2 (U-test, *P* = 0.0022) Fig.1E, filled brown circles).

Previous studies have reported that TBS-induced LTP is unable to support the expression of STC (*16, 17*). To verify this, STC was examined in rat hippocampal slices (Fig. 1F). TBS-LTP was induced in input S1 (filled blue circles), and 30 min later WTET was delivered to S2 (open blue circles) to induce early-LTP. Consistent with earlier findings, no expression of STC was observed. Potentiation in S1 remained statistically significant for up to 6 h (Wilcox test, *P* = 0.0156), whereas in S2 late-LTP was not expressed and synaptic responses decayed to baseline within 200 min (Wilcox test, *P* = 0.0625).

Given that rodent hippocampal slices fail to express STC during TBS-induced activity, we next asked whether TBS-LTP in NHP hippocampal slices can engage STC. After a stable 30-min baseline recording, TBS was applied to input S1 (Fig. 1G, filled brown circles). Thirty minutes later, early-LTP was induced in S2 by WTET (Fig. 1G, open brown circles). Strikingly, early-LTP in S2 was transformed into late-LTP, indicating robust expression of STC. Both S1 and S2 exhibited significant potentiation immediately following TBS and remained potentiated throughout the entire 6-h recording period (Wilcox test, *P* = 0.0313 for S1; Wilcox test, *P* = 0.0313 for S2 at 360 min post-tetanization).

Together, these findings demonstrate that stable and long-lasting LTP can be induced in NHP hippocampal slices and, importantly, that TBS-LTP in NHPs supports STC. This suggests that NHP hippocampal circuits may possess a distinct proteomic environment that facilitates the synaptic tagging and capture of plasticity-related proteins.

### Divergent Gene Expression and Proteomic Signatures in Rat and Non-Human Primate Hippocampi

It is well established that BDNF, CaMKII, CaMKIV, PKMζ, and CREB play critical roles in the maintenance of long-term synaptic plasticity in the hippocampal CA1 region of rodents (*3*). We therefore assessed mRNA expression levels in rat and NHP hippocampal slices under control conditions and following TBS-induced LTP (Fig.2). In NHP hippocampal slices, TBS-LTP resulted in a significant increase in BDNF and CaMKIV mRNA expression in the CA1 region compared with control slices without TBS. In contrast, the expression levels of CaMKII, PKMζ, and CREB did not show significant changes (Fig.2A). Interestingly, TBS did not induce significant changes in the expression of any of the candidate molecules in rat hippocampal slices (Fig. 2B). In NHP hippocampal slices, TBS-LTP induced a significant increase in BDNF (t test, *P* = 0.0234) CaMKIV (t test, *P* = 0.0362) mRNA expression in the CA1 region compared with control slices without TBS (Fig. 2A). In contrast, the expression levels of CaMKII (*P* = 0.0786), PKMζ (*P* = 0.3657), and CREB1 (*P* = 0.5700) were not significantly changed. In rat hippocampal slices, TBS did not produce significant changes in the expression of any of the examined molecules (Fig. 2B), including BDNF (*P* = 0.2495), CaMKII (*P* = 0.9195), CaMKIV (*P* = 0.5554), PKMζ (*P* = 0.9407), and CREB1 (*P* = 0.2794).

**Figure 2.**
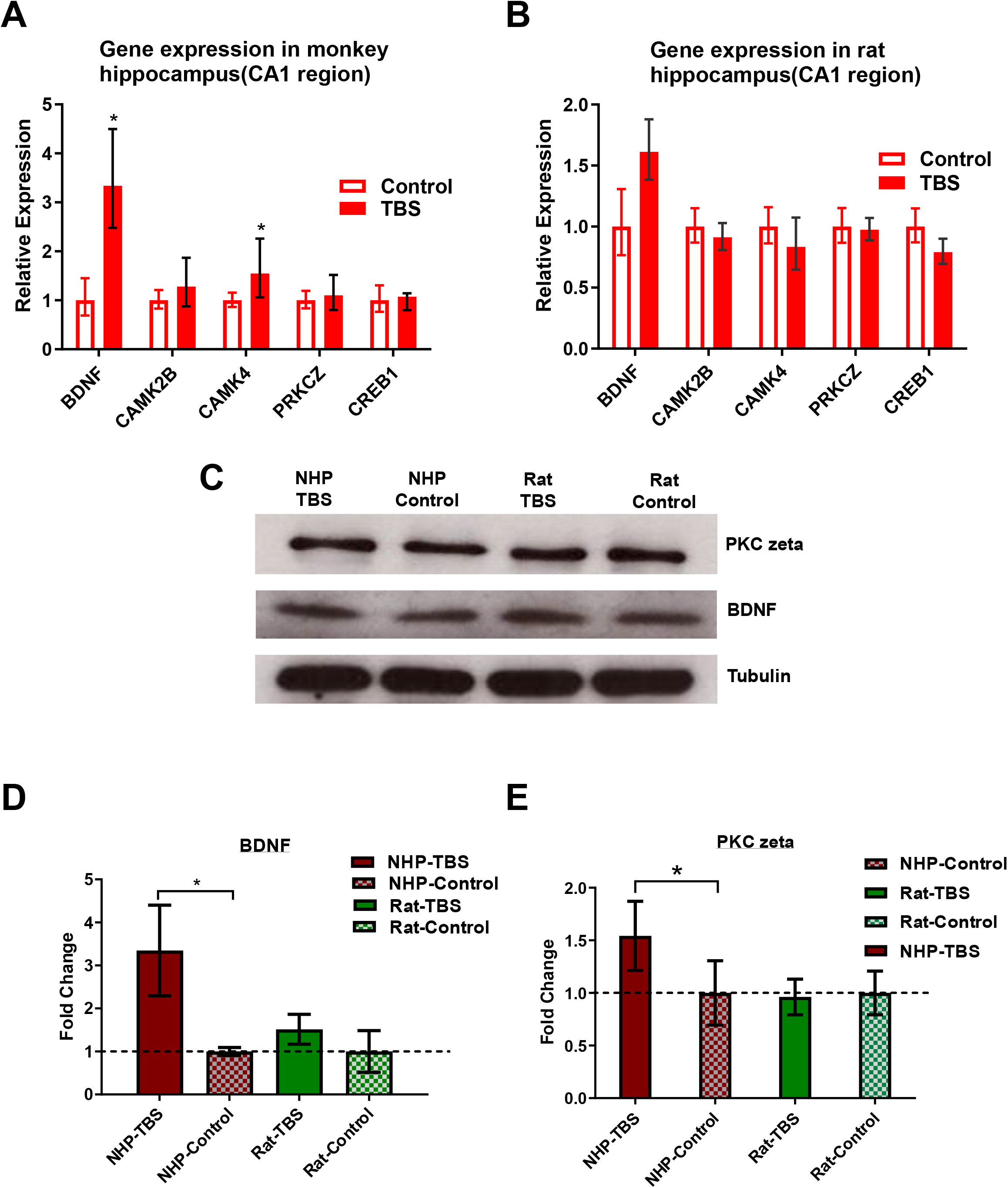
Divergent gene expression and proteomic signatures in rat and nonhuman primate (NHP) hippocampal CA1 following TBS-induced LTP. **(A-B)** Relative mRNA expression levels of plasticity-related genes in hippocampal CA1 under control conditions (black) and following TBS-induced LTP (red). (A) In NHP hippocampal slices, TBS significantly increased the expression of *BDNF* and *CaMKIV*, while *CaMKII, PKMζ*, and *CREB1* remained unchanged. (B) In rat hippocampal slices, TBS did not induce significant changes in the expression of any of the examined genes. **(C)** Representative Western blot images showing protein levels of PKMζ and BDNF in NHP and rat hippocampal slices under control conditions and following TBS. Tubulin was used as a loading control. **(D-E)** Quantification of protein expression levels. (D) BDNF and (E) PKMζ protein levels were significantly increased following TBS in NHP hippocampal slices compared to controls, whereas no significant changes were observed in rat hippocampal slices.

Given the established roles of PKMζ and BDNF in maintaining long-term plasticity and synaptic tagging and capture in hippocampal CA1 (*16*), we next examined their protein expression levels in rat and NHP hippocampi following TBS. TBS induced a significant increase in both PKMζ (t test, *P* = 0.0221) and BDNF (t test, *P* = 0.0184) protein levels in NHP hippocampus but not in rat hippocampus (Fig. 2C-D-E). These results suggest that TBS-induced up-regulation of PKMζ in the NHP hippocampus may contribute to the enhanced STC observed in NHPs compared with rodents.

### The Roles of BDNF and PKMζ in LTP maintenance in rats and NHPs

It has been previously reported that blocking PKMζ 1 h after tetanization reverses established LTP (*22, 23*), whereas inhibition of PKMζ after tetanization fails to disrupt potentiation in DHPG-primed TBS-LTP (*16*). We first examined whether BDNF contributes to TBS-LTP in rat hippocampal slices. To this end, recombinant human TrkB/Fc chimera (TrkB/Fc; 1 μg/mL) was applied 60 min after the induction of TBS-LTP and maintained until the end of the experiment (Fig.3A). TrkB/Fc scavenges extracellular BDNF and thereby blocks TrkB activation. Blockade of BDNF impaired the late phase of LTP (Fig. 3A, filled blue circles), resulting in a decaying potentiation that persisted only up to 250 min relative to its own baseline (Wilcoxon test, *P* = 0.0313) and until 250 min when compared with S2 (U-test, *P* = 0.0238).

**Figure 3.**
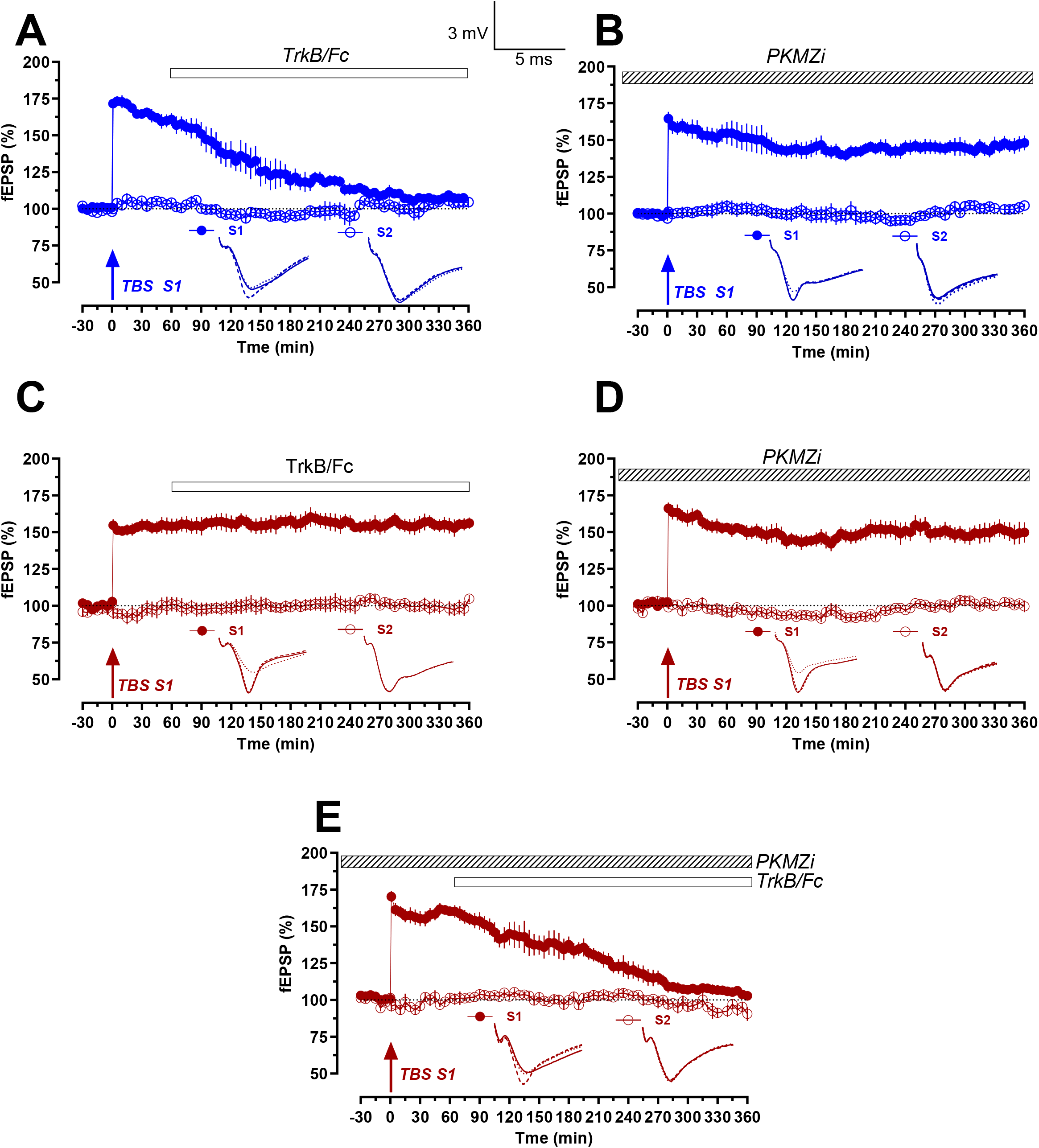
Differential roles of BDNF and PKMζ in the maintenance of TBS-induced LTP in rat and nonhuman primate (NHP) hippocampal CA1. **(A-B)** Rat hippocampal slices. (A) Blockade of BDNF signaling using TrkB/Fc applied 60 min after TBS impaired the late phase of LTP, resulting in a gradual decay of potentiation in S1, while the control pathway S2 remained stable. (B) In contrast, inhibition of PKMζ (PKMZi), applied prior to TBS and maintained throughout the recording, did not affect LTP maintenance, indicating that PKMζ is not required for TBS-induced LTP in rat CA1 and that it can be maintained exclusively by BDNF. **(C-D)** NHP hippocampal slices. (C) Inhibition of BDNF signaling with TrkB/Fc did not disrupt TBS-induced LTP, with S1 responses remaining persistently potentiated throughout the recording period. (D) Similarly, inhibition of PKMζ alone did not affect LTP maintenance, indicating that TBS-induced LTP in NHP hippocampus is resistant to disruption of either pathway individually. Asterisks denote statistically significant differences between groups (*P < 0.05). **(E)** Combined inhibition of BDNF and PKMζ in NHP hippocampal slices. Simultaneous blockade of both pathways prevented the maintenance of TBS-induced LTP, resulting in a decay of potentiation in S1 toward baseline levels, while S2 remained stable. This indicates cooperative and partially redundant roles of BDNF and PKMζ in sustaining long-term synaptic plasticity in NHP CA1. In all panels, figures shown in blue represent experiments conducted in rats, whereas those in dark red represent experiments in nonhuman primates (NHPs). Solid arrow indicate the time points of application of TBS. A hatched or empty rectangle represents the timing and duration of application of specific inhibitors. Analog traces depict representative S1 and S2 fEPSPs: 30 min before any experimental manipulation (dotted line), 30 min (hatched line) and 360 min (solid line) after TBS or WTET. Vertical scale bar: vertical: 3 mV, horizontal scale bar: 5 ms.

We next assessed the role of BDNF in NHP TBS-LTP by applying TrkB/Fc (1 μg/mL) 60 min after TBS-LTP induction and continuing the application until the end of the recording (Fig. 3C). Surprisingly, BDNF blockade had no significant effect on LTP maintenance in NHP hippocampal slices (Fig. 3C, filled brown circles), resembling the BDNF independence previously reported for DHPG-primed TBS-LTP (*16*). Potentiation in S1 was statistically significant immediately after TBS and remained stable throughout the entire 6-h recording period (Wilcoxon test, *P* = 0.0313; U-test, *P* = 0.0022 at 360 min post-tetanization). We then investigated the role of PKMζ in TBS-induced LTP in both rat and NHP hippocampal slices. Application of a PKMζ inhibitor (PKMZi) 60 min before TBS-LTP induction and maintained throughout the remaining recording period had no effect on TBS-LTP maintenance in either species (Fig. 3B and 3D). In both cases, LTP persistence after TBS remained significantly elevated relative to the corresponding baseline and the independent control input S2 (Wilcoxon test, *P* = 0.0313; U-test, *P* = 0.0022 for rat; Wilcoxon test, *P* = 0.0313; U-test, *P* = 0.0022 for NHP at 360 min post-tetanization).

Thus, whereas inhibition of BDNF selectively disrupts the late phase of TBS-LTP in rat hippocampal slices while sparing PKMζ-dependent mechanisms, TBS-LTP in NHP slices is resistant to inhibition of either BDNF or PKMζ alone. We therefore tested the combined effect of BDNF and PKMζ inhibition (Fig. 3E). Strikingly, simultaneous inhibition of BDNF and PKMζ prevented the maintenance of TBS-LTP in NHP hippocampal slices (filled brown circles), with statistically significant potentiation lasting only up to 245 min relative to its own baseline (Wilcoxon test, *P* = 0.0313) and up to 270 min when compared with the independent control input S2 (U-test, *P* = 0.0411).

Together, these findings indicate that TBS-LTP in NHP hippocampus is supported by cooperative and partially redundant molecular mechanisms that differ fundamentally from those operating in rodent hippocampus.

## Discussion

Our findings demonstrate that TBS induces persistent LTP in both rodent and NHP hippocampal slices. TBS closely mimics endogenous theta-frequency firing patterns associated with exploratory behavior and memory encoding and is therefore widely used as a physiologically relevant model of synaptic plasticity (*24, 25*). In rodents, TBS-induced LTP is largely localized, translation-dependent, and relatively independent of transcription, resulting in spatially restricted availability of plasticity-related proteins (PRPs) (*26-29*). This compartmentalization limits its ability to support synaptic tagging and capture (STC), which requires broader PRP availability (*16*).

In contrast, we observed that TBS-LTP in NHP hippocampus facilitates STC, indicating a fundamental divergence in the mechanisms underlying LTP persistence across species. Importantly, the threshold for inducing short-lasting LTP via weak tetanization was comparable between rodents and NHPs, suggesting that baseline synaptic excitability is conserved. Therefore, the observed differences are unlikely to arise from altered induction thresholds but instead reflect divergence in downstream molecular processes governing PRP availability and synaptic stabilization.

A key finding of this study is the differential recruitment of PRPs following TBS. In rodents, TBS-LTP critically depends on BDNF signaling, as BDNF sequestration abolishes the expression of TBS-LTP (*16, 27*). Consistent with this, inhibition of PKMζ does not affect LTP maintenance in rats, suggesting that PKMζ is not recruited following TBS-induced potentiation (*16, 23*). In contrast, our data show that TBS in NHP hippocampus leads to increased expression of both BDNF and PKMζ. Given that PKMζ is a key determinant of PRP availability and synaptic stabilization, its recruitment in NHPs may lower the threshold for STC and facilitate persistent plasticity. This is consistent with previous observations in primed LTP paradigms, such as DHPG-facilitated TBS-LTP, where enhanced PRP synthesis supports long-lasting potentiation (*16*). However, simultaneous inhibition of both pathways abolished LTP maintenance, indicating that BDNF and PKMζ may function in a compensatory or cooperative manner in primates. This redundancy may enhance the robustness and flexibility of synaptic plasticity in species with more complex cognitive capabilities.

Molecular analyses further revealed species-specific differences in gene and protein expression profiles following TBS. In NHPs, elevated mRNA levels of BDNF and CaMKIV indicate a greater contribution of transcription-dependent mechanisms to LTP maintenance, whereas such changes are not observed in rodents under TBS. At the protein level, elevated PKCζ and BDNF in NHPs further support enhanced PRP availability. These findings are consistent with previous studies demonstrating divergence in the expression and regulation of PRPs between rodents and primates, despite conservation of core synaptic machinery (*21*). Beyond molecular mechanisms, structural and circuit-level differences between rodent and primate brains-including divergence in synaptic organization, connectivity, and neuronal diversity-may also contribute to the observed effects (*21, 30*). Variations in neuronal density, synaptic organization, and interneuron diversity can influence network dynamics and integration properties (*31, 32*). Additionally, primate-specific transcriptional programs and epigenetic regulatory mechanisms may modulate activity-dependent gene expression, including plasticity-related genes such as Arc, Egr1, and c-Fos (*33, 34*). These differences may extend the temporal window for PRP availability and enhance the efficiency of synaptic tagging and capture; however, this remains to be investigated, particularly in light of recent rodent evidence suggesting that tag–PRP interactions can persist longer than originally proposed (*35*).

Collectively, our findings support the view that STC represents a conserved framework for memory-related plasticity, but that the molecular pathways governing PRP production and utilization have undergone species-specific specialization. In rodents, TBS-LTP relies predominantly on BDNF-dependent mechanisms, whereas in primates, additional recruitment of PKMζ and transcriptional pathways enhances the probability of long-term synaptic stabilization.

In conclusion, our study demonstrates that physiologically patterned stimulation engages distinct molecular programs in the rodent and NHP hippocampus, leading to differences in the associativity of synaptic plasticity. These findings reveal fundamental differences in the molecular architecture underlying plasticity across species and underscore the importance of incorporating NHP models in translational neuroscience research. Reliance solely on rodent systems may overlook critical regulatory mechanisms relevant to human cognition and memory disorders. Therefore, understanding species-specific molecular cascades, particularly those governing STC, will be essential for translating synaptic plasticity mechanisms to human cognition and for developing effective therapeutic strategies targeting memory disorders.

## Materials and Methods

### Preparation of hippocampal slices

A total of 36 acute hippocampal slices were obtained from 7 male Macaca fascicularis (5-7years old), and 31 hippocampal slices were obtained from 15 male Wistar rats (5-7 weeks old). All animal procedures were approved by the Institutional Animal Care and Use Committee (IACUC) of the National University of Singapore.

Briefly, rodents were decapitated after anesthetization using CO_2_. NHPs were euthanized using pentobarbital (150 mg/kg). Adequate euthanasia was confirmed by cyanosis of the mucous membranes and the absence of palpebral reflex, heart rate, and respiratory movements., In both cases, the brains were quickly removed and cooled in 4°C artificial cerebrospinal fluid (ACSF) that contained the following (in mM): 124 NaCl, 3.7 KCl, 1.0 MgSO_4_ .7H_2_O, 2.5 CaCl_2_, 1.2 KH_2_PO_4_, 24.6 NaHCO_3_ and 10 D-glucose, equilibrated with 95% O_2_–5% CO_2_ (carbogen; total consumption 16 L/hr), and acute hippocampal slices were prepared from the right hippocampus using a manual tissue chopper. Hippocampal slices were then transferred onto the interface brain slice chamber (Scientific Systems Design) and incubated for three hours at 32°C with ACSF before the electrophysiology studies In all electrophysiological recordings, two-pathway experiments were conducted. Two monopolar lacquer-coated stainless steel electrodes (5 MΩ; AM Systems, Sequim) were positioned at an appropriate distance within the stratum radiatum of the CA1 region to stimulate two independent synaptic inputs, S1 and S2, within a single neuronal population. These electrodes elicited field excitatory postsynaptic potentials (fEPSPs) from Schaffer collateral/commissural-CA1 synapses (Figure 1a). Pathway specificity was confirmed following the method described by (*16*). A third electrode (5 MΩ; AM Systems) was placed in the apical dendritic layer of the CA1 region to record fEPSP. The signals were amplified using a differential amplifier (Model 1700; AM Systems), digitized with a CED 1401 analog-to-digital converter (Cambridge Electronic Design), and monitored online. After the pre-incubation period, a synaptic input-output curve (afferent stimulation vs. fEPSP slope) was generated. The test stimulation intensity was adjusted to evoke an fEPSP slope corresponding to 40% of the maximal response for both S1 and S2 synaptic inputs. To maximize the efficient use of NHP hippocampal tissue, electrophysiological recordings were performed using three interface chambers, each accommodating two slices for simultaneous recordings. Distinct stimulation paradigms were applied concurrently across chambers, enabling multiple experimental conditions to be examined in parallel within the same biological preparation. This approach allowed systematic comparison of experimental manipulations while minimizing the number of animals required.

Late long-term potentiation (L-LTP) was induced using a theta-burst stimulation (TBS) protocol similar to that described previously (*16*). The protocol consisted of 50 bursts, each containing four stimuli delivered at an interstimulus interval of 10 ms. The bursts were applied at 5 Hz (inter-burst interval of 200 ms) over a period of 10 s (*16*). For early LTP induction, a “weak” tetanization (WTET) protocol comprising a single stimulus train of 21 pulses at 100 Hz (0.2 ms stimulus duration per polarity) was used, as described by (*36*). In all experiments, a stable baseline was recorded for at least 30 minutes using four biphasic constant-current pulses (0.1 ms per polarity) delivered at 0.2 Hz at each time point.

### Pharmacology

The PKMζ antisense oligodeoxynucleotide (IDT, Singapore) was stored as a 2 mM stock solution in Tris-EDTA (TE, pH 8.0) buffer at −20 °C (Tsokas et al., 2016). Stock solutions were stored for no longer than one week. Before use, the stock was diluted to the required final concentration in artificial cerebrospinal fluid (ACSF), bubbled with carbogen (95% O_2_ / 5% CO_2_), and bath-applied for the specified duration. The final concentration of PKMζ antisense oligodeoxynucleotide used in the experiments was 20 μM.

TrkB-Fc chimeric protein (Cat. #688-TK, R&D Systems, Minneapolis, MN, USA) was prepared fresh for each experiment by dissolving the lyophilized protein in distilled water (dH_2_O) to obtain a final concentration of 1 μg/mL.

### mRNA Quantitative Real-Time PCR

For mRNA expression analysis, cDNA synthesis was performed using the GoScript Reverse Transcription System (Promega, USA). Briefly, 2 μg of RNA was preheated with 2 μL of Oligo (dT) at 72°C for 2 minutes. Reverse transcription was conducted at 42°C for 1 hour, followed by enzyme inactivation at 95°C for 5 minutes. Quantitative real-time PCR (qRT-PCR) was carried out using the StepOne Plus Real-Time PCR System (Applied Biosystems) with TaqMan Universal PCR Master Mix (Cat. No. 4304437; Thermo Scientific) and TaqMan probes specific for BDNF, CaMKII, CaMKIV, PKMζ and CREB. The qRT-PCR protocol involved an initial denaturation step at 95°C for 10 minutes, followed by 40 amplification cycles of 95°C for 15 seconds and 60°C for 1 min. The fold changes in BDNF, CaMKII, CaMKIV, PKMζ and CREB. gene expression were calculated using the 2^−^ΔΔCt method (Livak & Schmittgen, 2001), with GAPDH serving as the internal normalization control. Each sample was measured in duplicate, and gene expression analysis was performed on hippocampal slices obtained from three independent biological variants in each group (rodent or NHP; n = 3).

### Western blot analysis

In brief, hippocampal slices from four experimental groups (NHP TBS-LTP, NHP control, WT TBS-LTP, and WT control) were collected 4 h after TBS stimulation, flash-frozen in liquid nitrogen, and stored at −80 °C. Total protein was extracted using TPER Tissue Protein Extraction Kit (Thermo Fisher Scientific) supplemented with Halt Protease Inhibitor Cocktail (Thermo Fisher Scientific). Protein concentrations were determined by Bradford assay (Bio-Rad). Equal amounts of protein (20 µg per sample) were resolved by 10% SDS/PAGE and transferred to PVDF membranes. Membranes were incubated overnight at 4 °C with primary antibodies against CREB (1:500; Cell Signaling Technology) or BDNF (1:1,000; Abcam), followed by appropriate secondary antibodies. α-Tubulin (Sigma-Aldrich) was used as a loading control. Immunoreactive bands were visualized using SuperSignal West Pico Plus Chemiluminescent Substrate (Pierce Biotechnology) and quantified with ImageJ. Band intensities were normalized to the corresponding α-tubulin signal. For all experiments, slices obtained from a minimum of three independent biological samples (n= 3) were used for analysis.

### Data Analysis and Statistics

In field electrophysiological recordings, the strength of synaptic responses was quantified by measuring the slope of the fEPSP (millivolts per millisecond). All data are presented as mean ± SEM. To assess statistical significance within groups, the Wilcoxon signed-rank test (denoted as Wilcox) was applied to compare the mean normalized fEPSP values at specific time points with the fEPSP at −15 minutes (baseline). For comparisons between different groups, the Mann-Whitney U test (denoted as U test) was used. Statistical significance was set at *p < 0*.*05*. Nonparametric tests were chosen because a Gaussian normal distribution could not always be assumed, especially given the small sample size and the analysis of prolonged recordings (*37, 38*). In vitro electrophysiology data are presented as “n,” representing the number of slices.

For qRT-PCR and Western blot analyses, data from three to four independent experiments are presented as mean ± SD. Statistical significance was determined using Student’s t test or one-way ANOVA, as appropriate. Differences were considered significant at *P* < 0.05. Analyses were performed using Prism (GraphPad Software).

## Author Contributions

Conceptualization: SS and CL; Methodology: AM,KK,YSC, WLW,SN, YPW,STW and SS; Investigation: AM,KK,YSC, WLW,SN; Visualization: AM,KK,YSC, WLW,SN; Funding acquisition: SS and CL; Project administration: SS and CL; Supervision: SS and CL; Writing – original draft: AM, SN and SS; Writing – review & editing: SN, WLW, CL and SS.

## Acknowledgments

This work was supported by the NUHS Seed Fund (NUHSRO/2024/089/T1/Seed-Mar24/02) and the Ministry of Health (MOH-000641-00 and MOH-001883-00) awarded to S.S. C. L was supported by MOE-T2EP30121-0010.

